# Mycorrhizal dominance reduces forest-tree species diversity

**DOI:** 10.1101/2021.01.23.427902

**Authors:** Alexis Carteron, Mark Vellend, Etienne Laliberté

## Abstract

Ectomycorrhizas and arbuscular mycorrhizas, the two most widespread plant-fungal symbioses, are thought to differentially influence tree species diversity, with positive plant-soil feedbacks favoring locally abundant ectomycorrhizal tree species and negative feedbacks promoting species coexistence and diversity in arbuscular mycorrhizal forests. While seedling recruitment studies and cross-biome patterns of plant diversity and mycorrhizal dominance support this hypothesis, it remains to be tested at the forest stand level over continental scales. Here, we analyze ∼85,000 forest plots across the United States to show that both ectomycorrhizal-dominated and arbuscular mycorrhizal-dominated forests show relatively low tree diversity, while forests with a mixture of mycorrhizal strategies support a higher number of tree species. Our findings suggest that mycorrhizal dominance, rather than mycorrhizal type, shapes tree diversity in forests.

## Introduction

Mycorrhizas – the most widespread terrestrial symbiosis on Earth – have long been known for their nutritional benefits to plants^1^. However, there is increasing interest in their role as drivers of local plant biodiversity^2^. Species-rich tropical rainforests are mainly composed of arbuscular mycorrhizal (AM) trees^3^ while species-poor boreal forests are dominated by ectomycorrhizal (EcM) trees^4,5^ suggesting that the AM strategy favors plant species coexistence and diversity while the EcM strategy promotes dominance by one or few species^2,6^. Small-scale studies of seedling recruitment support this hypothesis: EcM seedlings perform better when growing in soils near (or conditioned by) conspecific individuals (i.e., showing positive plant-soil feedbacks), whereas the opposite has been found for AM plants^7,8^. Proposed mechanisms for positive feedback in EcM forests include greater protection to conspecific seedlings from soil-borne pathogens and improved nutrient acquisition, relative to AM forests^6,9^. However, we do not know whether these short-term effects on recruitment dynamics translate into persistent effects on canopy tree species composition and diversity. Indeed, neither the historical biome-level observations nor the individual-level studies of seedling recruitment directly test the hypothesis that EcM-dominated forests sustain lower tree species diversity than AM-dominated forests; broad-scale analyses at the forest-tree community level are needed to resolve this.

Here, we use an extensive grid-based inventory of 84,448 naturally forested plots surveyed by the U.S. Department of Agriculture Forest Service (Forest Inventory and Analysis program) to explore the relationship between mycorrhizal dominance (EcM vs. AM strategy) and local tree species diversity across broad environmental gradients at the continental scale. Selected plots (each composed of four subplots of 168 m^2^) spanned the contiguous U.S.^10^, with the number of tree species per plot ranging from 1 to 21 (Fig. 1A). Mycorrhizal strategy for the 357 tree species present in the selected plots was extracted from a recent published database^11^. As a predictor of tree diversity at the plot scale, we calculated the proportion of total basal area, from stem diameter measurements, comprised of trees with the same mycorrhizal strategy. Because the vast majority of plots are dominated by EcM and/or AM strategies (Fig. S1), patterns of EcM and AM proportions are essentially mirror images (Fig. 1B, Fig. S2). We hypothesized that tree diversity would decrease monotonically as dominance by EcM trees increased, such that AM forests would show the highest diversity (see “the ectomycorrhizal dominance hypothesis”, Fig. 2A). We tested for a relationship between tree diversity and EcM proportion in several ways: (i) using the simple bivariate relationship (results in Supplementary Material); (ii) after controlling for other environmental factors; and (iii) after controlling for the statistical necessity that forests with multiple mycorrhizal types have a greater species pool of trees to draw from than forests with just one mycorrhizal type.

**Figure 1.**
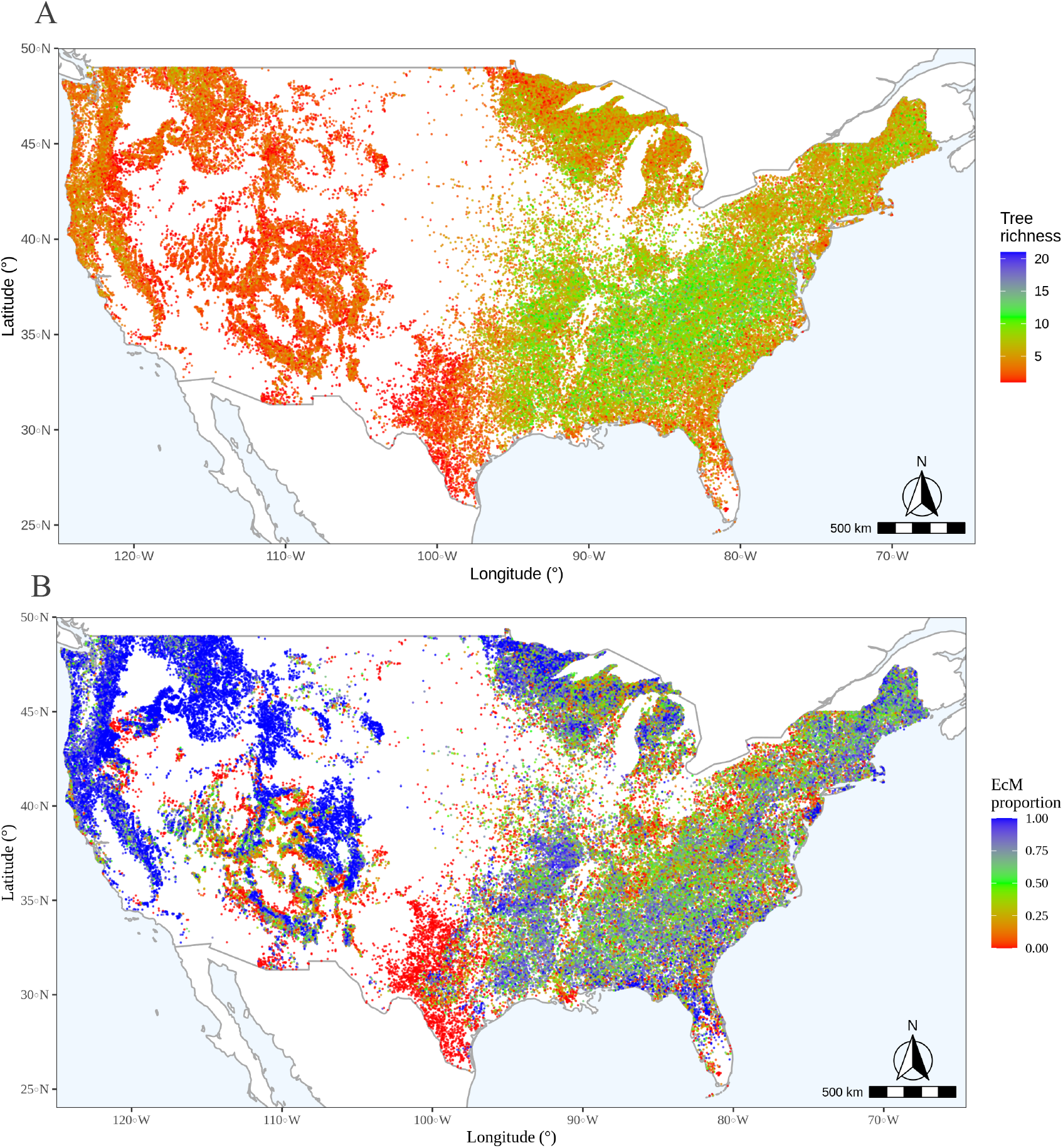
Tree richness and ectomycorrhizal (EcM) proportion at the community scale across the U.S. (A) Map of tree richness (number of tree species); (B) Map of EcM proportion (as the proportion of basal area per plot of trees with DBH > 12.7 cm known to associate with EcM fungi).

**Figure 2.**
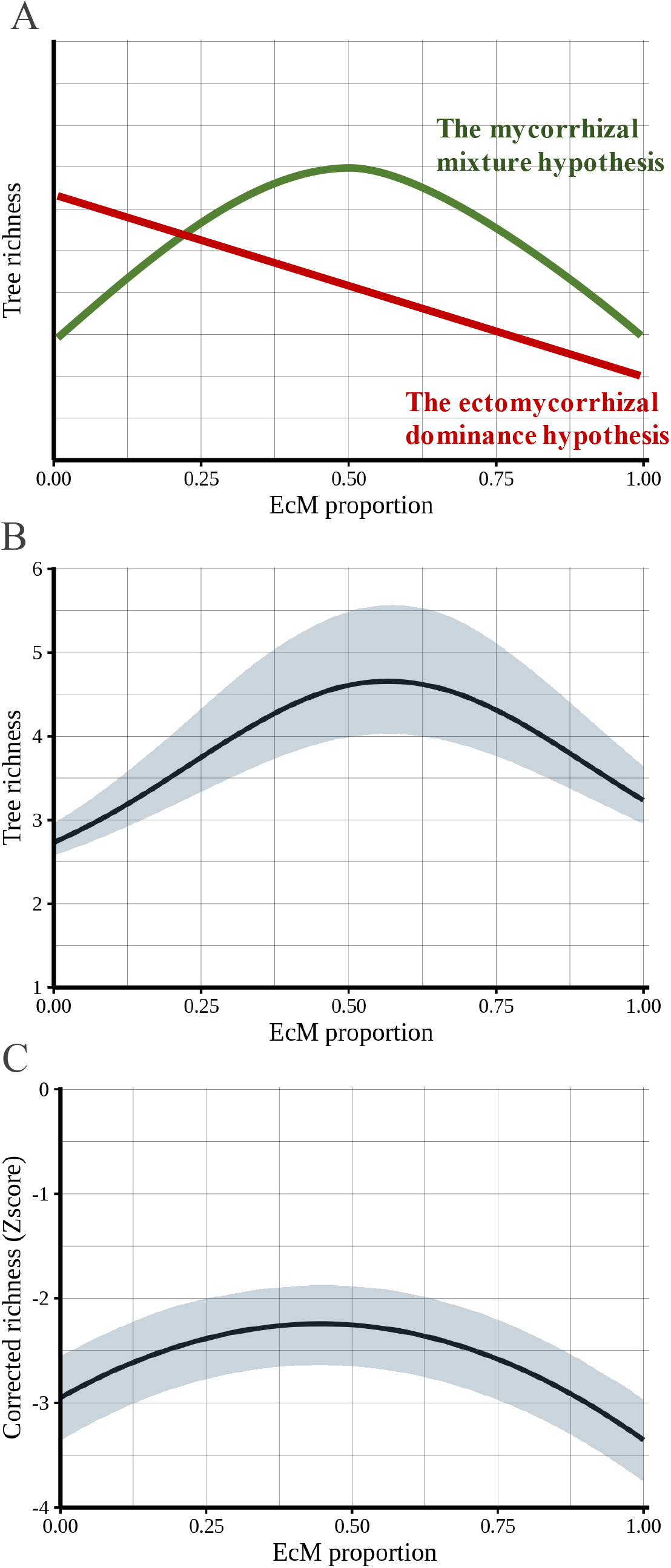
Relationships between EcM proportion and tree richness. (A) Hypothetical relationships, (B) EcM proportion vs. predicted tree richness and (C) EcM proportion vs. corrected tree richness. Both (B) and (C) take into account potentially confounding environmental factors (elevation, orientation, physiography, precipitation, slope and temperature), and the lines indicate the regression curve between both variables and shaded areas represent 95% confidence interval of the regression.

## RESULTS

In line with our hypothesis, the bivariate relationship showed that tree species diversity was relatively low in EcM-dominated forests (Fig. S3). However, contrary to our hypothesis, tree diversity was maximal when EcM tree basal area was ∼50%, and tree species diversity declined under increasing dominance by AM trees (Fig. S4; See “the mycorrhizal mixture hypothesis” in Fig. 2A). As such, tree species diversity was lowest in forests dominated by either the EcM or AM strategies, and highest when there was an approximate mixture of both strategies (Fig. S3, S4).

Mycorrhizal distributions are known to be correlated with environmental factors that also influence plant diversity^12^, but the pattern in the bivariate relationship remained strong after controlling for environmental variables (Fig. 2B). In models including effects of local abiotic factors (climatic, topographic and physiographic properties), we found that tree diversity was influenced by these factors, especially temperature, topography and water availability, but the negative effects of mycorrhizal dominance on tree diversity were strongest (Fig. 2B, Fig. 3).

**Figure 3.**
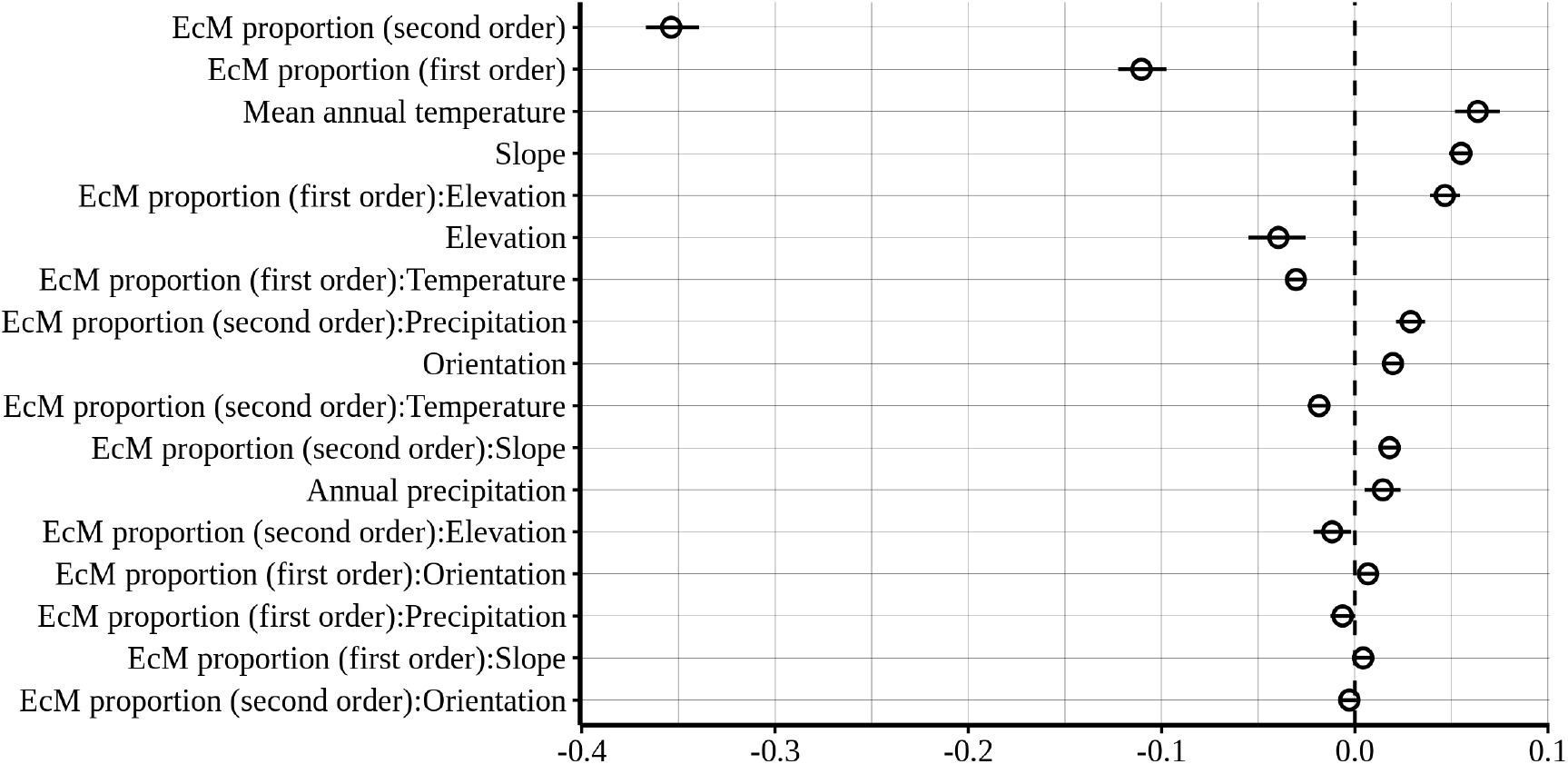
Posterior coefficient estimates. (medians are represented by empty circle and 95% credible intervals by black vertical line) for the ectomycorrhizal (EcM) proportion (first and second order terms) and local environmental factors (elevation, orientation, precipitation, slope, temperature), and their interactions on tree species richness. Variables were standardized prior modeling. Terms are ordered based on the absolute values of the slopes.

Our results are also robust to the potential statistical artefact that forest plots with only one mycorrhizal type have lower *potential* diversity (i.e., only tree species of that type can be present). Local plant diversity is determined by environmental filtering from the regional flora^13^, and species diversity depends on the size and composition of the regional species pool^14^. As such, we applied a null model to determine if mycorrhizal dominance had detectable effects on tree species diversity beyond what could be explained by the tree species composition of the regional flora. We assessed the expected relationship between tree diversity and the proportion of EcM tree basal area based on random sampling from the regional tree species pool, from which we calculated the deviation between observed and expected values for each plot (which we call “alpha-deviation”). The null model re-assigned a species identity to each individual tree in a given plot based on a random draw of species of the same mycorrhizal type (simplified as either EcM or “other” strategies) from the regional pool, with probability weights proportional to species regional abundances. Our null model thus preserved the value of EcM proportion in each plot. Regional pools were defined within each of 25 ecoregions^15^, which represent geographical areas with relatively similar ecological and environmental conditions (Fig. S5). Results of our null model analysis showed that the hump-shaped relationship with tree species diversity being lowest at low or high EcM basal area persisted (Fig. 2C, Fig. S6). This means that the lower local tree species diversity observed in plots dominated by either the EcM or AM strategy was not the result of the regional species pool containing a smaller number of tree species from either one of these strategies, but rather an outcome of local processes reducing species diversity in EcM- or AM-dominated plots.

## DISCUSSION

Our results strongly suggest that dominance by either EcM or AM strategy, and not only the EcM strategy type, reduces local tree diversity across the forested United States. Several mechanisms involving mycorrhizal type may combine locally to influence plant diversity^2^. First, positive plant-soil feedbacks commonly reported for EcM species at the seedling recruitment stages^7,8^, could also apply to AM species, but at later life stages (e.g. saplings or small sub-canopy trees), eventually leading to canopy dominance by certain AM tree species (Fig. S7). While it is widely accepted that EcM plants may benefit more from EcM fungi in terms of mineral nutrition and protection against root pathogens^9^, the higher maintenance costs of EcM fungi compared to AM fungi could mean that the net benefits of the two mycorrhizal types are similar^16^, thereby equalizing fitness differences among strategies. Furthermore, fine-scale niche partitioning could promote coexistence of different mycorrhizal types^17^ and ecosystems with a mixture of mycorrhizal strategies may also create environments that are more diverse and spatially heterogeneous^18^. Together, these processes could locally promote diversity where multiple plant nutrient-acquisition strategies such as mycorrhizal types co-occur^8,19^.

A number of studies have reported positive effects of a diverse inoculum of AM fungi on plant diversity and ecosystems functions^20,21^. Plant species richness as well as evenness increase in response to AM fungi inoculation^22^, and plant-soil feedbacks involving seedlings tend to be more negative in AM tree species^7^. These results lead to the expectation of a positive effect of AM dominance on tree diversity. However, studies at the community level are typically conducted in grasslands^23^, which may not apply to long-lived trees in forests. As a result, the few studies of trees are typically at the level of individual seedlings^7,8,24^, and these short-term effects on seedling recruitment might not necessarily translate to canopy-level patterns involving mature trees. It is also worth noting that even though most tropical rainforests are both AM-dominated and host high tree species diversity, there is also a number of hyper-dominant tropical tree species forming AM associations^25^. Therefore, not all tropical forest communities are species-rich, even in neo-tropical forests where AM trees dominate^26^. Furthermore, recent evidence had cast doubt on the conventional hypothesis of lower plant diversity in EcM compared to AM systems assessing temperate and boreal sites^27^. Using data at the continental scale, our results show that mycorrhizal dominance – regardless of mycorrhizal type – shapes tree species diversity in forests with diversity maximized when different mycorrhizal strategies co-exist.

In sum, patterns of mycorrhizal dominance and tree diversity have been historically considered among distant biomes, leading to the hypothesis that the EcM symbiosis reduces plant diversity, while the AM symbiosis promotes plant diversity^2,12^. The impact of mycorrhizal fungi on plant nutrition and plant soil-feedbacks, often studied at the individual level scale, have provided indirect support for these hypotheses^2^.

However, forest trees interact locally over prolonged periods, and results from our study based at the community and canopy-level suggest that short-term effects such as negative plant-soil feedback effects involving AM seedlings at the recruitment stage do not persist in the longer term to influence canopy tree species diversity. In fact, we find that mycorrhizal dominance (either AM or EcM) favors dominance by a single species, thus reducing diversity, and potentially leading to alternative stable states in forest tree species composition^28^. Even though mycorrhizal dominance can be determined at several scales (e.g., root system, stand, biome), our study highlights the importance of considering the impact of mycorrhizas on ecological processes at the scale of the forest plot (< 1 km^2^). At this scale, co-existing mycorrhizal strategies may act as a driver of plant diversity, which might only be detected by studying the entire gradient of mycorrhizal proportions. Forests with a mixture of mycorrhizal strategies are sometimes overlooked, as they are considered less common^4^. However, this is not the case across North America, and forests with diverse nutrient acquisition strategies may represent a crucial avenue for research and forest management targeting greater ecosystem services and adaptation to climate change.

## METHODS

### Data collection

For this study, we used publicly available data from the U.S. Department of Agriculture Forest Service, known as the Forest Inventory and Analysis (FIA) program. Data were accessed from https://apps.fs.usda.gov/fia/datamart/CSV/datamart_csv.html on 28 February 2020. The primary objective of the FIA is to determine the extent, condition, volume, growth, and use of trees on U.S. forest land in order to frame realistic forest policies and programs^10^. This database has been used to address many ecological questions across large scales and gradients^11,29,30^. Plots are distributed relatively evenly in forested areas across all of the lower 48 contiguous states. Plot location uncertainty is < 1.6 km^10^. Each standard plot consists of four 7.3-m radius circular subplots (168 m^2^) within which all stems > 12.7 cm diameter at breast height (DBH) are identified to species and measured. There is one center subplot surrounded by the three peripheral subplots, each at a distance of 36.6 m from the center subplot.

Prior to our analyses, the data set was filtered using several criteria following the FIA user guide for Phase 2^10^. We only kept: (i) census data from the most recent year available for each plot, (ii) standard production and standardized plots (i.e. “Sample kind code” of 1, 2 or 3) that were sampled the same way (i.e. “Plot design code” of 1, 220, 240, 311, 314, 328, 502, 505) (iii) data taken using the National Field procedures, (iv) forested, natural and undisturbed stands with no observable recent silvicultural treatment. If data were missing for any measured values or variables the plots were excluded. Otherwise, plots were retained if more than four individual trees were present. In total, we analyzed data for 84,448 plots containing 2,518,123 trees.

For each selected plot, topographic data (elevation, slope, aspect) and physiographic class (estimate of moisture available to trees) were accessed directly from the FIA database. Climatic data (i.e. average annual temperature, annual precipitation) were accessed from the PRISM (Parameter-elevation Regressions on Independent Slopes Model) climate group (800-m spatial resolution; available at http://prism.oregonstate.edu/).

From the stem diameter measurements, total basal area was calculated for each species in each plot. Mycorrhizal strategy for each tree species was determined using data from Jo *et al*.^11^. Species were listed as either ectomycorrhizal (EcM), arbuscular mycorrhiza (AM), non-mycorrhizal (NM) or both AM and EcM (AM+EcM).

Tree species richness was calculated as the number of observed tree species with DBH > 12.7 cm. Species richness (Fig. 1A) followed large scale tree diversity gradients previously mapped for the U.S.^31^. Abundance was incorporated into diversity indices using the exponential of Shannon’s entropy index (*q* = 1) and the inverse of Simpson’s concentration index (*q* = 2), calculated as proposed in Chao *et al*.^32^.

Following the “National hierarchical framework of ecological units”^15^ we defined 25 ecoregions and assigned one for each plot depending on its location (Fig. S5). Ecological units are defined as areas of similar surficial geology, lithology, geomorphic processes, soil groups and subregional climate.

### Null model

We used a null model that re-assigned the species identity of the individuals in each plot based on random draws from the regional pool of tree species within ecoregions (Fig. S5), while keeping the total number of individuals per plot and the proportion EcM constant. Each species’ abundance (i.e., its probability of being chosen by the null model) was calculated as the number of tree stems in the ecoregion, divided by the total number of stems across species. We ran 100 randomizations from which we calculated the diversity “deviation” (or “corrected” diversity) as the observed diversity minus the mean of the null distribution of diversity values, divided by the standard deviation of this distribution. Diversity measures were the same as for the observed data. Negative values of corrected diversity represent lower diversity than expected given random draws from the regional species pool, which can be the result of environmental, demographic and stochastic processes that exclude some species locally. The null model was implemented in R^33^ and the code is available at [provide DOI upon publication].

### Modeling

To quantify the effect of mycorrhizal proportion and the environmental variables on tree species diversity and corrected diversity, we used generalized linear mixed-effect models implemented in a Bayesian framework. Ecoregion was included as a random factor. For richness values (*q* = 0), we used the Poisson distribution and for *q* = 1 and *q* = 2 we used the Gamma distribution. Because diversity values start at one, distributions were truncated with lower bound < 1. For the corrected diversity, we used the Gaussian distribution. Before modeling, variables were scaled by. subtracting the mean and dividing by the standard deviation.

Analyses on diversity were conducted for *q* = 1 and *q* = 2, which showed similar patterns as *q* = 0 (Fig. S8). Robustness to the minimum number of trees per plot of the relationship between tree richness and EcM proportion was tested by running models with a threshold for a minimum number of individuals per plot of nine and 14 individuals (Fig. S9). They showed similar pattern with a slight increase in maximum tree richness with an increasing threshold.

The models ran on four parallel chains of length 5,000 with a burn-in of 1,000 iterations with a thinning rate of 10. Uninformative priors were used as provided in the *brms* package^34^. Convergence was assessed for each parameter estimate by visually inspecting the Markov chains and considered sufficient when the 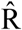 was equal to one.

Data manipulation and statistical analyses were done using the R software ^33^ and the following main packages: *brms*^34^, *data*.*table*^35^ *dplyr*^36^, *ggplot2*^37^, *ggpubr*^38^, *ggspatial*^39^, *raster*^40^, *reshape*^41^, *sf*^42^, *tidyr*^43^, *vegan*^44^. Scripts for all the data manipulation and analyses can be found at [provide DOI upon publication].

## ACKNOWLEDGMENTS

Funding, including scholarships to AC, was provided by Discovery Grants to EL (RGPIN-2014-06106, RGPIN-2019-04537) by the Natural Sciences and Engineering Research Council of Canada (NSERC) as well as a “Nouveau Chercheur” grant (2016-NC-188823) by the Fonds de recherche du Québec sur la Nature et technologies (FRQNT). AC would like to sincerely thank the following institutions for providing generous scholarships: FRQNT (Dossier 272522) and Université de Montréal through the “Bourse d’excellence Hydro-Québec”

## AUTHOR CONTRIBUTIONS

EL, AC and MV conceived the ideas and designed the methodology. AC, EL and MV analyzed the data and interpreted the results. AC led the writing of the manuscript. All authors contributed critically to the drafts and gave final approval for publication.

## COMPETING INTERESTS

Authors declare no competing interests.

## DATA AND MATERIALS AVAILABILITY

Data are available at https://apps.fs.usda.gov and the code is available at [provide DOI upon publication].

## Notes

### Competing Interest Statement

The authors have declared no competing interest.

