## Supplementary information for "Mycorrhizal dominance reduces forest-tree species diversity"

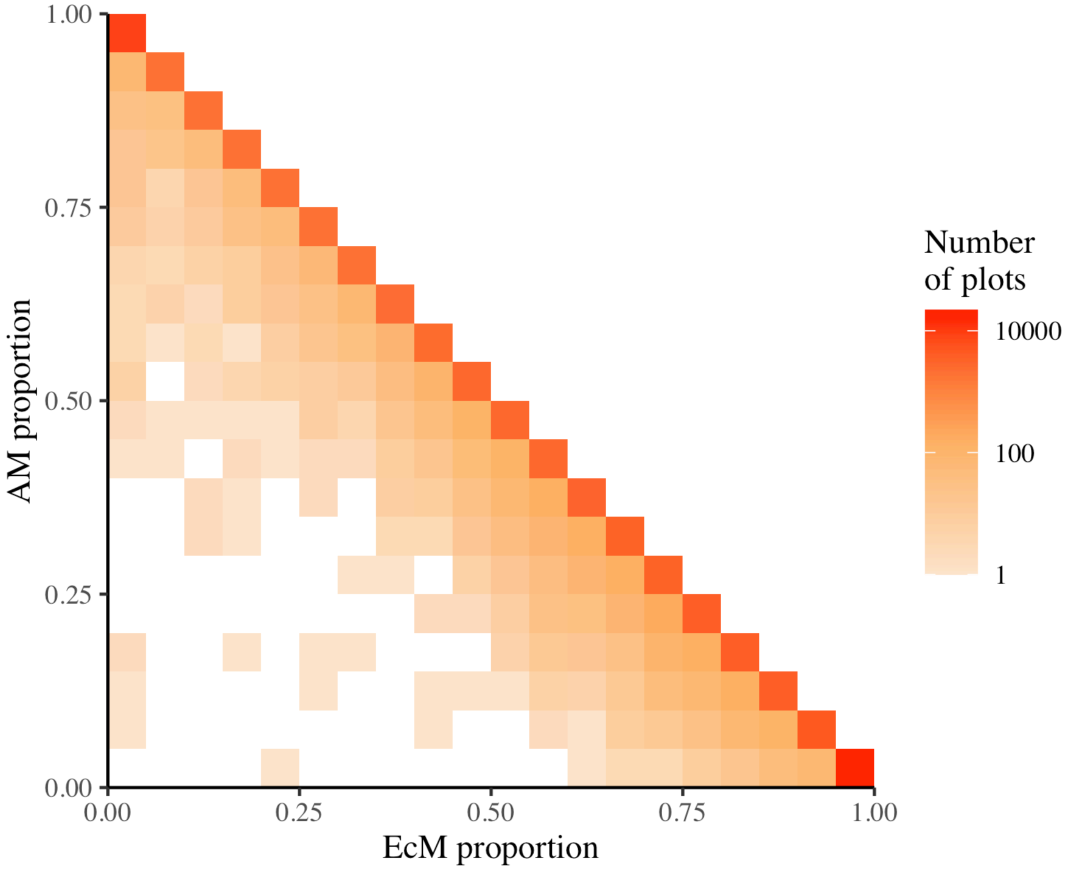


**Figure S1.** Relationship between ectomycorrhizal (EcM) and arbuscular mycorrhizal (AM) proportions in each plot. 95 % of the plots have a cumulative sum of AM and EcM proportions >.99 (i.e. most plots are located on the diagonal). The number of plots in the legend is presented on a log scale.

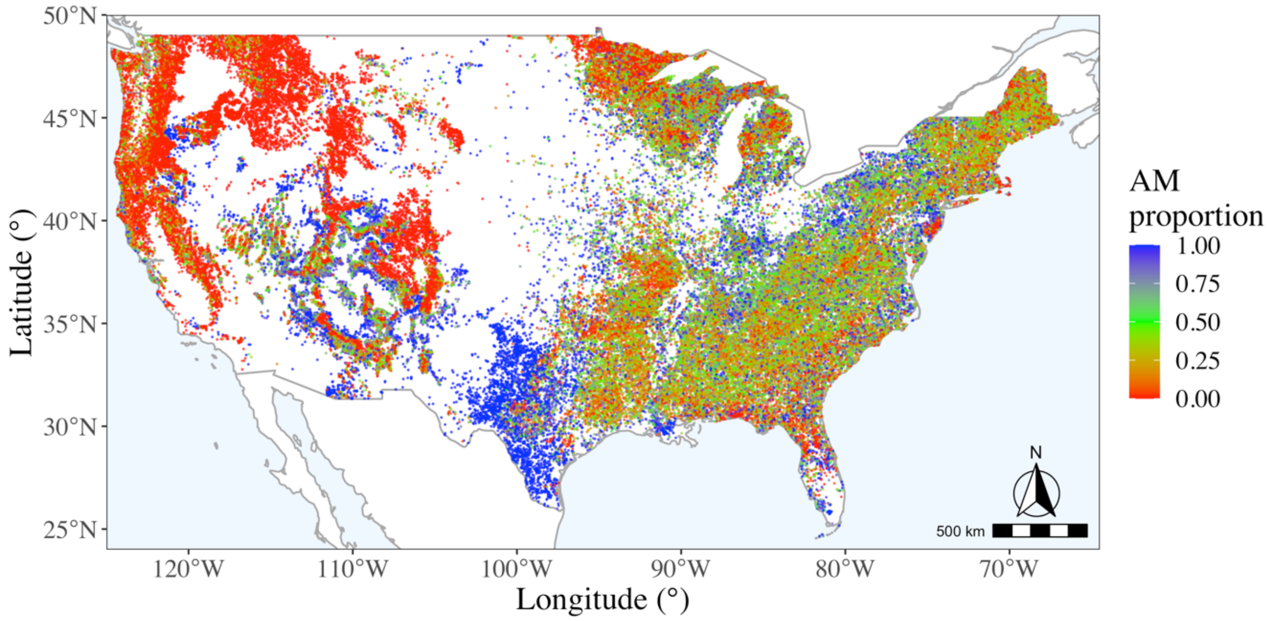


**Figure S2.** Map of arbuscular mycorrhizal (AM) proportion (as the proportion of basal area per plot of tree with DBH > 12.7 cm known to associates with AM fungi).

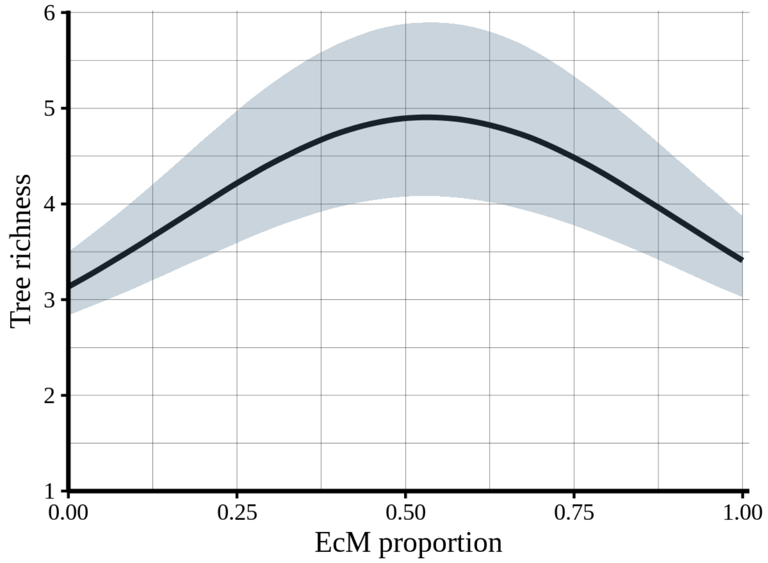


**Figure S3.** Relationship between EcM proportion and predicted tree richness without taking into account the other environmental factors. The line indicates the regression curve between both variables and the shaded area represents 95% confidence interval of the regression.

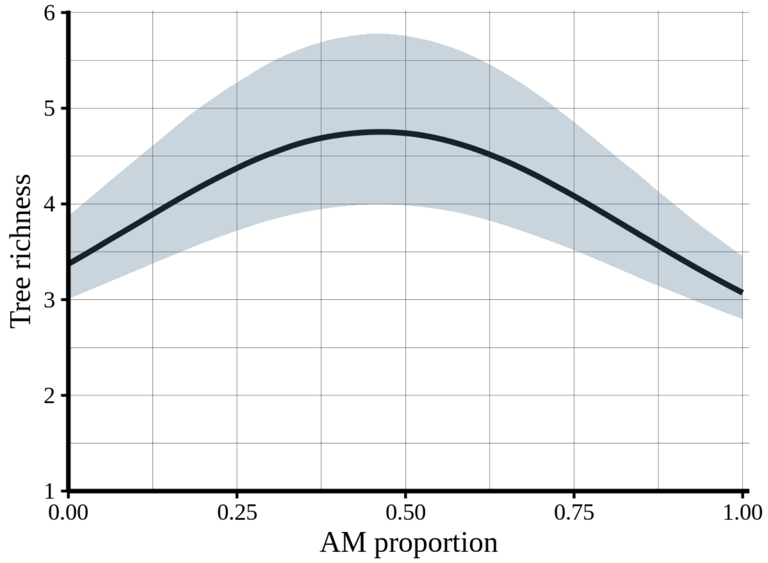


**Figure S4.** Raw relationship between AM proportion and predicted tree richness without taking into account the other environmental factors. The line indicates the regression curve between both variables and the shaded area represents 95% confidence interval of the regression.

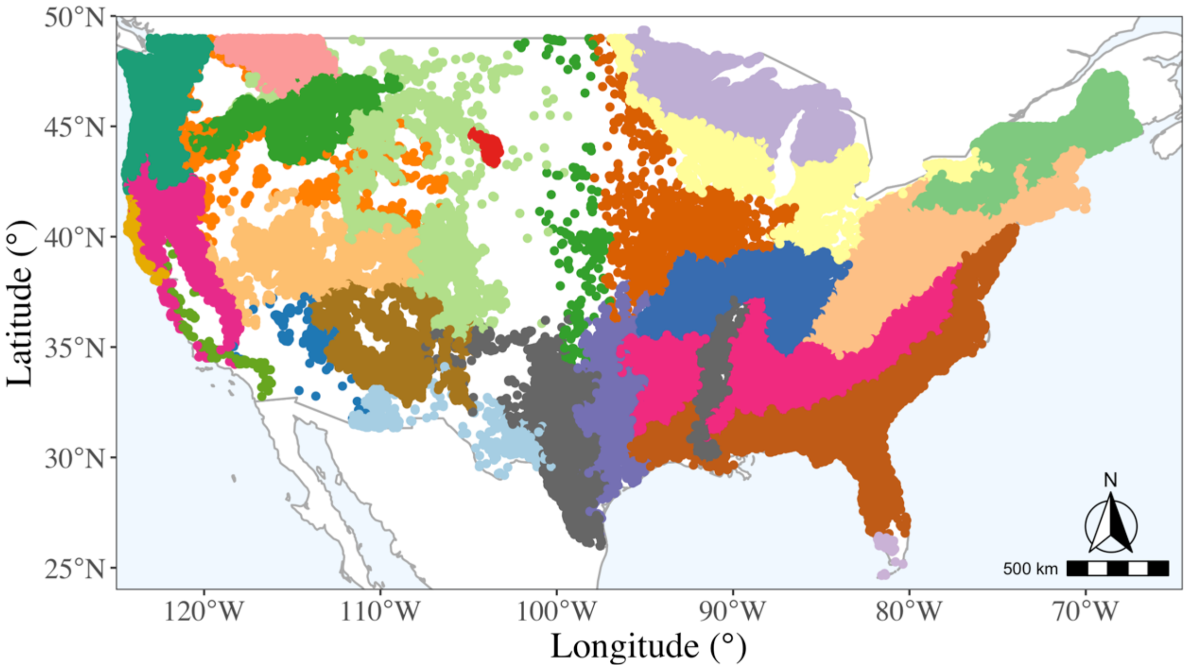


**Figure S5.** Map showing the plot location in the 25 ecoregions used in the models as random factor and in the null models as regional pools. Based on the "National hierarchical framework of ecological units" (*7*).

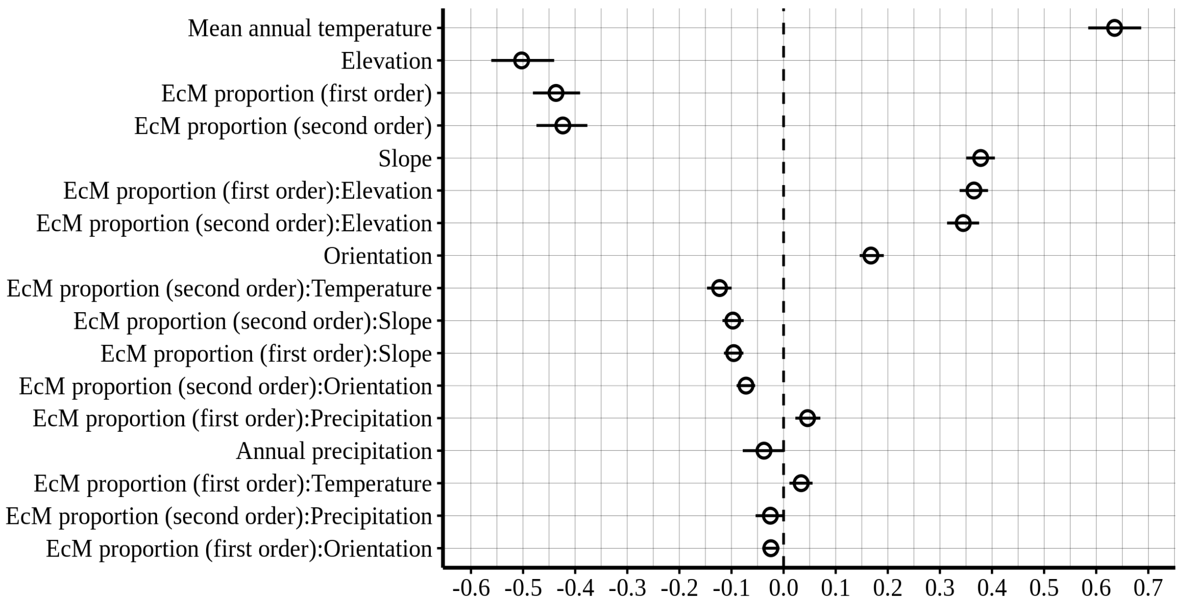


**Figure S6.** Posterior coefficient estimates (medians are represented by empty circle and 95% credible intervals by black vertical line) for the ectomycorrhizal (EcM) proportion (first and second order terms) and local environmental factors (elevation, orientation, precipitation, slope, temperature), and their interactions on alpha diversity deviation of tree species. Variables were standardized prior modeling. Terms are ordered based on the absolute values of the slopes.

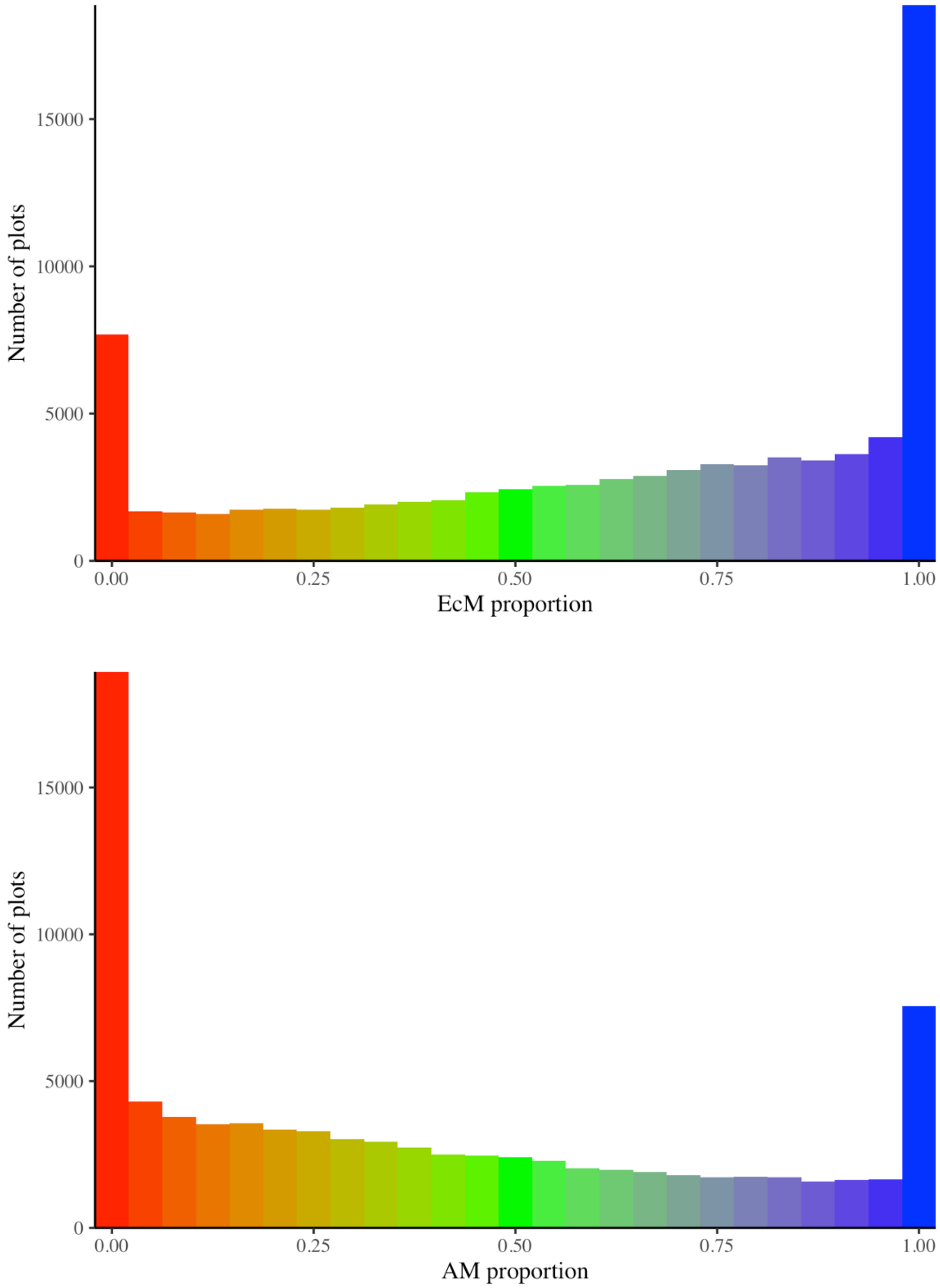


**Figure S7.** Relationships between the number of plots and ectomycorrhizal (EcM) proportion (top), and arbuscular mycorrhizal (AM) proportions (bottom).

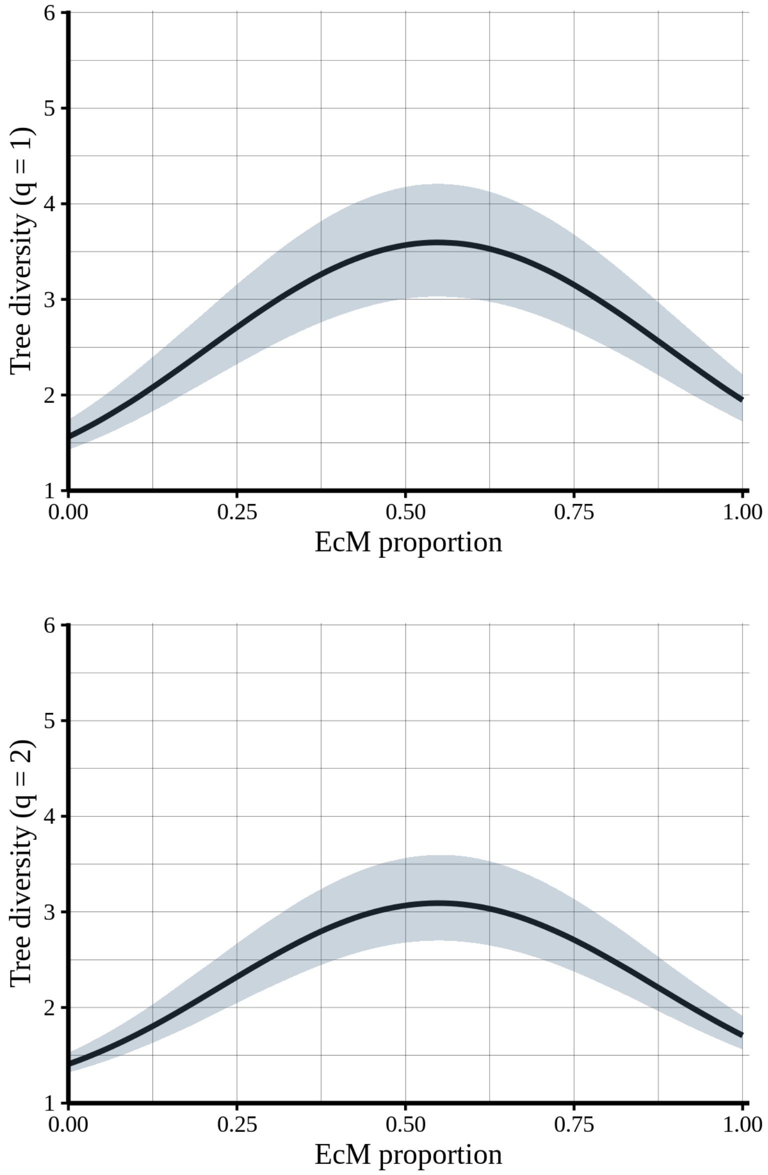


**Figure S8.** Relationships between ectomycorrhizal (EcM) proportion and model-predicted values of diversity for $q$ = 1 (top) and $q$ = 2 (bottom). Lines indicates regression curve between both variables. Shaded areas represent 95% confidence interval of the regression.

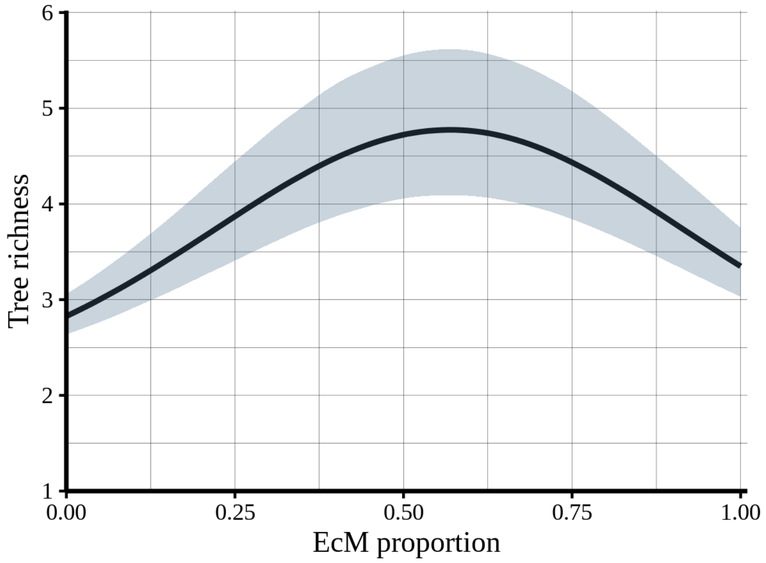


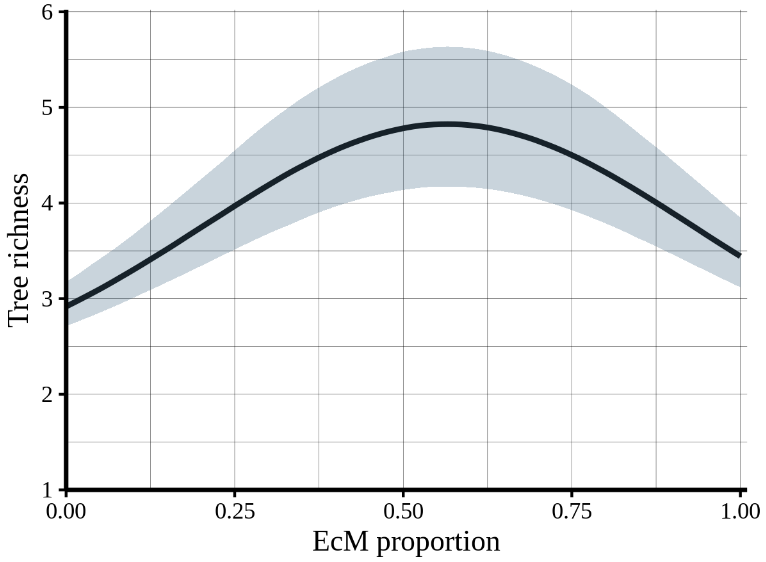


**Figure S9.** Relationship between EcM proportion and tree richness using thresholds for minimum number of individuals per plot of nine (top) and 14 (bottom) individuals. Lines indicates regression curve between both variables. Shaded areas represent 95% confidence interval of the regression.
